# Hypothesis: A plastically-produced phenotype predicts host specialization and can precede subsequent mutations in bacteriophage

**DOI:** 10.1101/411983

**Authors:** Colin S. Maxwell

## Abstract

The role of phenotypic plasticity in the evolution of new traits is controversial due to a lack of direct evidence. Phage host-range becomes plastic in the presence of restriction-modification (R-M) systems in their hosts. I modeled the evolution of phage host-range in the presence of R-M systems. The model makes two main predictions. First, that offspring of the first phage to gain a new methylation pattern by infecting a new host make up a disproportionate fraction of the subsequent specialist population, indicating that the plastically-produced phenotype is highly predictive of evolutionary outcome. Second, that the first phage gain this pattern is not always genetically distinct from other phages in the population. Taken together, these results suggest that plasticity could play a causal role on par with mutation during the evolution of phage host range. This uniquely tractable system could enable the first direct test of ‘plasticity first’ evolution.

## Introduction

Phenotypic plasticity is ubiquitous in nature but its role in evolution is controversial. The ‘plasticity first’ hypothesis holds that environmentally induced phenotypes frequently precede genetic changes during the evolution of new traits (1,2). Following the initial induction of the plastically-produced phenotype by the environment, this hypothesis holds that selection could ‘fix’ (make non-plastic) the trait through genetic assimilation (3), or refine the organism’s phenotype through genetic accommodation (1). Some even argue that plasticity fundamentally alters the logic of evolution by allowing non-genetic events to causally influence its outcome (4). Others doubt that genetic assimilation (5,6) or other varieties of plasticity-first evolution are common enough in nature to justify such a conclusion, or argue that whatever role plasticity plays in evolution can be understood without such a fundamental rethinking (7). This controversy persists because there are no systems where the causal role of plastically-produced phenotypes can be directly tested.

Directly testing whether plasticity causally influences the evolution of a trait requires testing if a plastically-produced phenotype both predicts evolutionary outcome (which individuals produce descendants with an evolved trait) and precedes any subsequent mutations that affect the trait. This is a complementary approach to comparative studies (2,8,9) or proofs-of-principle using artificial selection (10,11). However, it would require observing individuals in a population from the time when environmental conditions initially produced a phenotype via plasticity until a trait of interest evolved (12), which is impossible in almost all circumstances. Nevertheless, evolution can occur rapidly (reviewed in (13)), and several instances of new traits and even incipient species have been observed (14,15). Therefore, a strategy to resolve to this conundrum is to find systems where plasticity should play an important role *a priori* in the evolution of some trait and then observe the evolution of that trait in the laboratory by natural selection. By establishing a tractable system, a causal role for plasticity could be tested. Furthermore, since laboratory evolution can be replicated, the factors that determine *to what degree* a plastically-produced phenotype predicts evolutionary outcome and *how often* the phenotype precedes subsequent mutations could be determined, which could shed light on patterns of evolution outside the laboratory.

Experimental evolution using viruses that infect bacteria (bacteriophages or phages) is a powerful system for studying the evolution of new traits because phages have short generation times and high mutation rates. The types of bacteria a phage strain can infect (the host range) is a critical phenotype that determines both its niche and which other phage it can exchange genes with (16). This makes phage host range an excellent experimental model to test the origin of new traits.

Phage host range can be decomposed into a ‘genetic’ and ‘plastic’ component when the bacterial host has a restriction-modification (R-M) system. Phage have proteins that bind to host receptors that contribute to the genetic basis of host-range. An important class of host-range mutations are those affecting proteins that bind to host receptors (17-19). Since binding is determined by the sequences of the phage gene and the bacterial receptor gene, when the temperature and chemical composition of their surroundings is held constant, the component of host range caused by these proteins is ‘genetic’. Conversely, bacterial R-M systems can cause a plastic component in phage host range, as explained below. R-M systems are ubiquitous in prokaryotes (20) and have long been thought to protect their hosts from mobile genetic elements such as phage and plasmids (21). These systems encode restriction endonucleases, which cleave DNA at particular sites, and methyltransferases, which modify DNA at those sites (22). Genomic DNA is protected from cleavage by the restriction endonuclease through the activity of the methyltransferase, whereas invading DNA is recognized by the restriction endonuclease and cleaved before it can parasitize the cell.

If a phage evades the R-M system of a new host by chance (odds vary between 1 in 10 to 1 in 10 million (23)) and successfully infects it, that phage’s offspring’s fitness on the new host is plastically increased. This is because some fraction of progeny resulting from such infections will be marked with the methylation pattern of the new host by its methyltransferase and will therefore be invisible to that R-M system during subsequent infections. This fraction (the ‘methylation efficiency’) can vary between ∼100% for phage lambda (24) and ∼10% for T7 (25). This methylation pattern is not inherited via factors encoded in the phage genome but is determined by the host. Since the phenotype (host range) of the phage is influenced by the environment that it was produced in, the host-range of phage can be plastic due to host R-M systems.

Although plasticity allows phage to exploit hosts with R-M systems, this plasticity can be costly. If the methylation efficiency of a host less than 100%, then offspring without the methylation pattern will have low fitness on any host with an R-M system. In this case, mutations affecting the recognition sites of R-M systems—which abolish both methylation and cleavage—can fix in the population (25). Indeed, genome-wide data show that sites recognized by R-M systems are avoided by at least some phage and bacteria (26), suggesting that this selective pressure is widely felt in bacteria and their parasites. Therefore, there are two ways to produce a phage capable of efficiently replicating in a host once it has injected its DNA into it: either plastically via methylation or genetically via mutations affecting the recognition sites of the R-M systems. Furthermore, since the methylation efficiency of hosts need not be 100%, the plastically produced host range phenotype can be less fit (‘costly’) relative to the genetically produced phenotype. ‘Costs of plasticity’ (reviewed in (27)) are thought to play an important role in providing the selective pressure to ‘fix’ (that is, make non-plastic) plastically produced phenotypes during genetic assimilation (3).

To summarize, plasticity has a large effect on phage fitness (increasing survival on the new host up to 10 million-fold (23)), and genomic evidence suggests that a cost of plasticity imposed by less than perfect methylation efficiency can shape phage genome evolution (26). Thus, the evolution of host-range in the presence of R-M systems is a premier system to test a causal role for plastically produced phenotypes on evolutionary outcome because short-term evolution could be linked to clade-level patterns of genome evolution.

I simulated a population of phages evolving in an environment containing two hosts with two distinct receptors and two distinct R-M systems. Under these conditions, I hypothesized that [1] the population of phages would evolve into two sub-populations specializing on one host each with distinct tail fiber affinities. Furthermore, I hypothesized that knowing which phages had the plastically produced host-range phenotype caused by the R-M system would [2] predict which phages would found this lineage of specialists, and [3] that this plastic phenotype could precede subsequent mutations in the tail fibers needed to specialize on that host. My simulations confirmed all three hypotheses, suggesting that phenotypic plasticity can play a similar role as mutation during the evolution of phage host-range. The metrics developed to quantify the effect of plasticity in the simulations could be used to test whether plastically-produced phenotypes play a causal role during the evolution of other traits.

## Methods

R-M systems create selective pressure to specialize for infecting only one species of bacteria because lineages of phage that efficiently bind to both species of bacteria lose a large number of their offspring when those offspring attempt to switch hosts. However, if a phage manages to infect the new host, the offspring of such phages find themselves on a reversed fitness landscape. Since adsorption rate and methylation pattern have an epistatic effect on fitness, previously disfavored mutations increasing binding to the new host become favored and vice versa (see supplementary results, Figure S1). I hypothesized that these offspring would evolve to specialize on the new host and competitively exclude the offspring of subsequent phage that breached the restriction barrier, therefore dominating the new host. To test if this scenario is plausible, I simulated phages evolving on a mixture of bacterial hosts with distinct R-M systems.

I examined a simple system with two species of bacteria (*A* and *B*) that differed in their R-M systems and a population of initially clonal phages marked with the methylation pattern of species *A*. I modeled two critical components of phage fitness: the affinity of phage tail fiber proteins for the receptors of bacteria, and the presence of an R-M system in the host. I simulated the evolution of the phage using an individual-based model—one which explicitly models the behavior of individuals. This approach is useful for examining the consequences of phenotypic plasticity because it allows the phenotype and genotype of an individual to be easily associated with the phenotype and genotype of its descendants. I implemented the model in Python (version 3.4) using the Mesa framework (https://github.com/projectmesa/mesa). I will briefly describe the model (see also Figure 1 for a graphical summary); for details, including a table of parameters, see the supplemental information.

**Figure 1.**
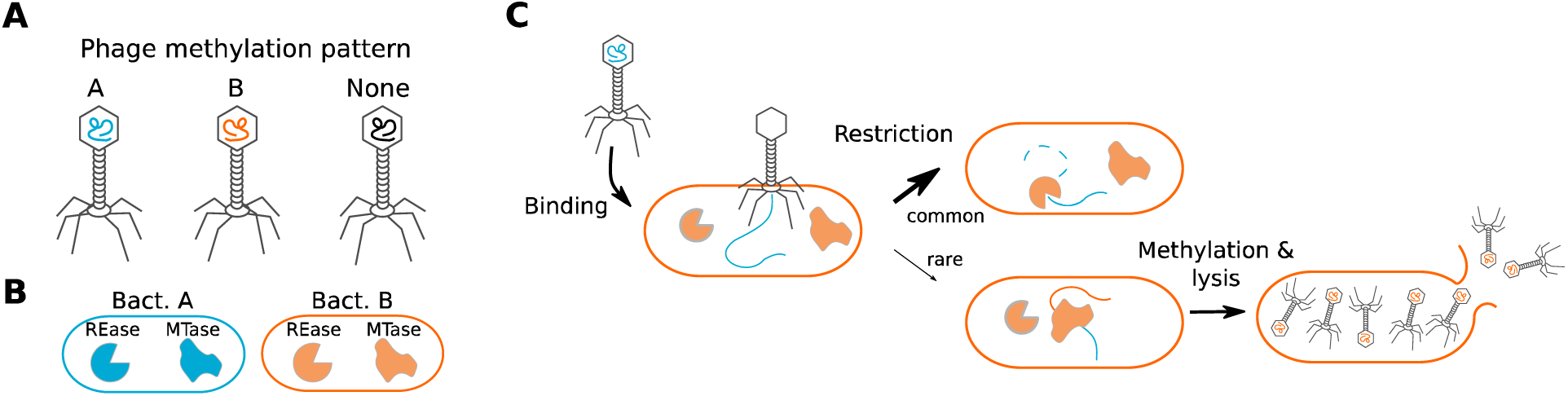
A schematic showing key elements of the model. **A** Bacteriophage were modeled as having DNA that can be methylated with either pattern ‘*A*’ or ‘*B*.’ **B** Bacteria were modeled as having distinct receptors and distinct restriction modification systems composed of a restriction endonuclease and a methyltransferase. **C** A schematic of the events that take place during each step of the simulation is shown. Phage bind to and inject their DNA into bacteria whereupon it is frequently degraded if the methylation pattern does not match the methylation pattern of the bacteria. If the phage is not killed by the R-M system, then it lyses the cell to produce progeny and the progeny is plastically marked with the methylation pattern of their host.

The phages evolved in a well-mixed environment constantly fed by bacteria without co-evolution between phage and bacteria. The number of bacteria was generally smaller than the equilibrium population of phages, indicating that there was competition for resources. During each time step in the model the phages were simulated encountering, binding to, injecting their DNA into, and producing progeny from bacteria. I modeled phages as having: [1] one of two methylation patterns, and [2] tail fibers that would bind to each bacterial species (Figure 1A) with different affinities (*p*_*A*_ and *p*_*B*_). Bacterial R-M systems destroyed DNA that was injected by a phage that was not marked with the cognate methylation pattern with some probability. Phage progeny genetically inherited their tail fiber affinity from their parent with mutation. I modeled five different ways for methylation to be produced: [1] randomly, [2] genetically, [3] 100% plastically, [4] 50% plastically, and [5] 10% plastically (Figure 1E). “Random” means phage get pattern *A* or *B* with 50:50 odds. “Genetic” means they inherit their methylation pattern from their parents with mutation. “*X%* Plastically” means that “*X%”* of phage have the methylation pattern of their host and the rest are unmarked. I did not model mutations affecting the recognition sites for the R-M system. For any given parameter set, I ran the simulation for 200 steps with 30 replicates.

The code used to generate all analyses is available at http://github.com/csmaxwell/phage-abm and is archived in Dryad (doi:TBD). The results of the simulations are archived in Dryad (doi:TBD).

## Results

### R-M systems select for host-range specialization

I first tested whether the simulated phage population would evolve specialist sub-populations that had affinity for only one bacterial species. I did not impose a trade-off between *p*_*A*_ and *p*_*B*_, so in the absence of an R-M system I expected generalists to evolve that would bind efficiently to both species (28). At the end of the simulation (200 generations), I examined *p*_*A*_ and *p*_*B*_ in individuals that had been produced from each species. Consistent with my expectations, phages only evolved specialist phenotypes when both restriction and non-random methylation were present (Figure 2). This indicates that in the presence of R-M systems, even with inefficient plasticity, phage evolve two distinct sub-populations of specialists.

**Figure 2.**
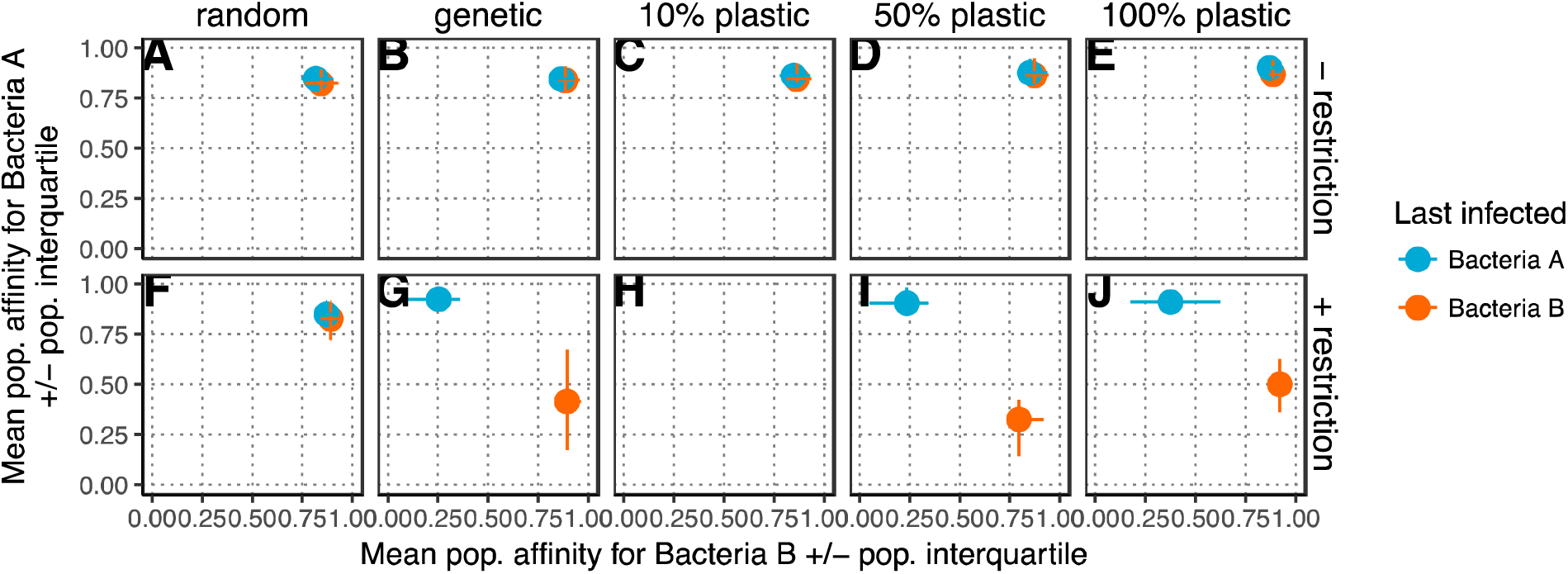
R-M systems select for host-range specialization. Plots of host-range specialization are shown for each methylation scheme in both the presence and absence of cleavage of improperly methylated DNA by bacteria. The average affinity for bacteria *A* and bacteria *B* (*p*_*A*_ and *p*_*B*_, respectively) of phage produced from bacteria *A* (blue) or bacteria *B* (orange) are shown at the end of the simulation (after 200 generations). The simulation was initialized with *p*_*A*_ = *p*_*B*_ = 0.5. The points show the average of the 30 replicates, error bars are bootstrapped 95% confidence intervals and are present on both X and Y axis, even when not visible. The rows of subplots show the results of the simulation run with (**A-E)** no restriction of improperly methylated DNA, or **(F-J)** with a restriction system with a 0.1% chance of restriction escape. The columns in the plot show the results of the simulation with **(A,F)** random methylation, **(B,G)** genetic inheritance of methylation, **(C,H)** 10% plasticity, **(D,I)** 50% plasticity, or **(E,J)** 100% plasticity in methylation. Missing points indicate that the phage population went extinct, which was only common when there was both restriction and 10% plastic methylation.

### The plastically produced phenotype predicts the pedigree of specialists

Each of the phages that make up the sub-population on the new host (*B*) must have come from lineages that breached the restriction barrier of that host at some point. At the end of the simulation (200 generations), there is a sub-population of specialist phages infecting the new host. How many lineages contribute to this population? I tested this by adding the number of phage equivalent to the progeny from one infection (0.1% of the starting population; the ‘test lineage’) at the beginning of the simulation, varied their methylation pattern and affinity for the new host, and recorded what percent of the specialist population *B* was derived from them. The plastically produced phenotype caused by breaching the restriction barrier is highly predictive of the pedigree of the specialist population—much more so than any mutation affecting tail fiber affinity (Figure 3). When restriction is present, mutations increasing *p*_*B*_ in the test lineage increased the fraction of phage derived from the test lineage in specialist population *B*, but only when they were marked with methylation pattern *B*. ‘Plastic’ methylation substantially increased the number of phages derived from the test lineage in population *B* relative to ‘random’ methylation. Notably, in simulations with both plastic methylation and restriction, a substantial proportion (∼50%-90%) of the phages infecting bacteria B were derived from the test lineage phages even when the test lineage phages had the same *p*_*B*_ as the rest of the population. This pattern held regardless of a trade-off between *p*_*A*_ and *p*_*B*_, and was robust to varying the simulation length, mutation frequency, and the efficiency of plasticity (Figures S1, S2).

**Figure 3.**
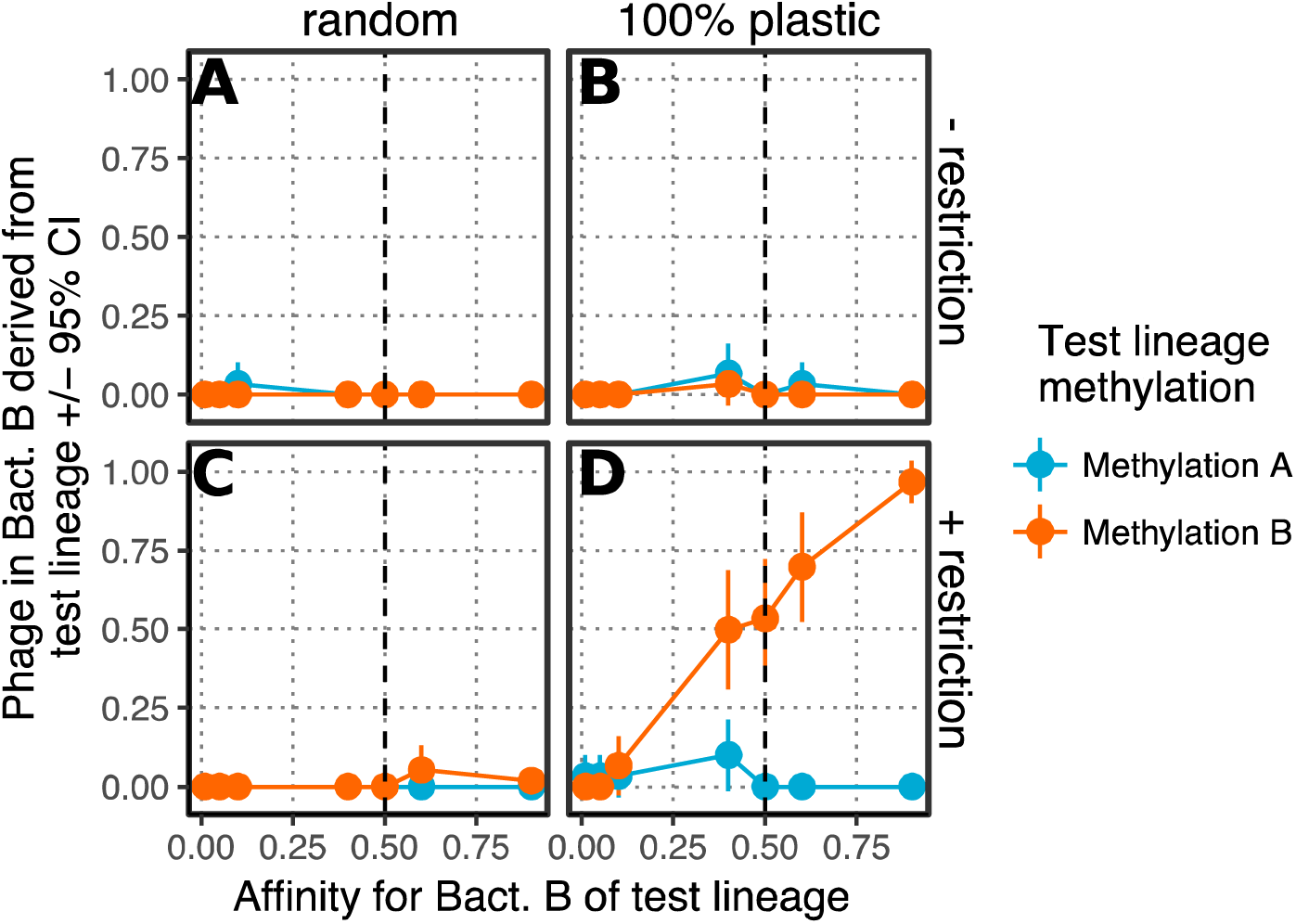
Knowing which phage first breached the restriction barrier of bacteria *B* predicts the pedigree of the bacteria *B* specialist population. The simulation was initialized with 1,000 phage with either *p*_*A*_ = *p*_*B*_ = 0.5 or *p*_*A*_ = 0.95, *p*_*B*_ = 0.05, and 10 phage (the ‘test lineage’) at the beginning of the experiment with different values of *p*_*B*_. The fraction of the phage infecting bacteria *B* at the end of the experiment (200 generations) that are derived from the test lineage phage is shown. The simulation was run with either **(A,B)** a 0.1% chance of restriction escape or **(C,D)** without restriction of improperly methylated DNA and with either **(A,C)** random or **(B,D)** 100% plastic methylation. Vertical dashed lines show *p*_*B*_ for the background population of phage at the start of the simulation. All simulations were run with no trade-off between *p*_*A*_ and *p*_*B*_. Error bars are 95% bootstrapped confidence intervals.

### Genetic diversity determines if methylation precedes mutation

Does the first phage to breach the restriction barrier during the simulations have a higher affinity for the new host than other individuals in the population? When the simulation was started with phages with some ability to bind to the new host or when mutation was rare, the affinity of the first phage to breach the barrier was similar to affinity for the new host in the rest of the population (Figure 4). However, when mutation was common or if the simulation was initialized with phage with no ability to bind to the new host, the first phage tended to be genetically distinct. This makes intuitive sense because the higher the pre-existing ability to bind to the new host, the less likely a mutation would be needed to allow binding. Calculations of the probability of infection and mutation confirmed that for realistic parameters of phage mutation rate and restriction bypass that a breaching the restriction barrier can precede mutation (Figure S3). When mutation is rare, R-M bypass is common, and when there is some pre-existing affinity for the new bacteria, then the plastically produced host range phenotype can precede mutation.

**Figure 4.**
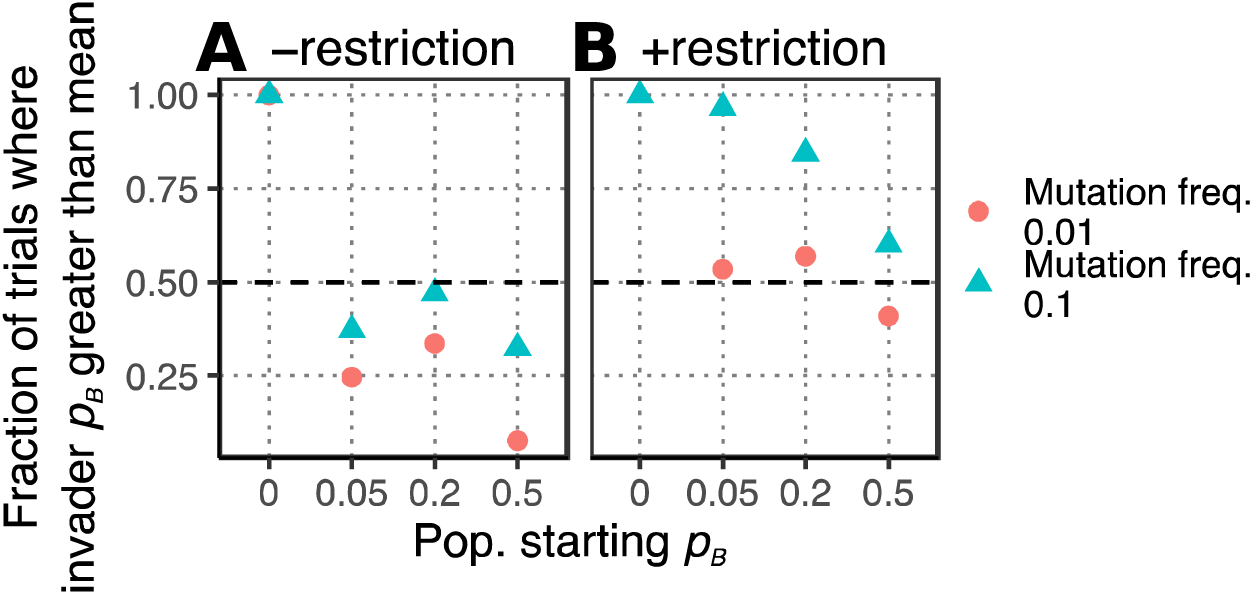
The order in which mutating to increase affinity for *B* and breaching the restriction barrier of *B* occurs in the individual-based model is determined by the frequency of mutation and the starting affinity of the population for *B*. The fraction of simulations where *p*_*B*_ of the first phage to successfully reproduce on *B* is greater than the mean *p*_*B*_ of the population is plotted. Simulations were run with either **(A)** no restriction of improperly methylated DNA, or **(B)** a 0.1% chance of restriction escape, either rare (pink dots) or common (teal triangles) mutation, and for various combinations of the population starting affinity for bacteria *B*. Only parameter sets where phage successfully reproduced in B in ten trials (out of 100 trials) are plotted.

## Discussion

The evolution of phage host specialization in the presence of R-M systems is an excellent system to examine the role of plasticity in evolution because host-range shifts can occur rapidly and reproducibly in the laboratory and because plasticity can have a large impact on the host-range of the phage. I used a simulation to explore how the plastic host range phenotype generated by R-M systems affects the evolution of host-range specialization. I used two metrics to measure how the plastically-produced phenotype affected phage evolution: ‘predictive power’ and ‘precedence.’ The host-range phenotype produced by the R-M systems *predicts* the pedigree of phages in the specialist population that evolves. Furthermore, since breaching the restriction barrier of a host can occur at a much higher rate than mutations in phage genomes, the plastic host-range phenotype can *precede* subsequent mutations needed to specialize on the new host. The model indicates that the plastic host-range phenotype can cause the evolution of a specialist population, but this prediction needs to be tested. The ability of phage to find and parasitize a new host is analogous to other examples of organisms encountering and then exploiting new niches, suggesting that when a plastically-produced phenotype has a large effect on the likelihood that an organism’s offspring will experience the same environment, that it could cause the evolution of specialists in these cases as well.

### Empirical predictions and possible tests

The model described here makes three main predictions. First, R-M systems impose a trade-off between the ability to exploit two hosts that leads to the evolution of host specificity. This could be tested using experimental evolution by serially passaging phage on strains that differed both by their receptors recognized by a phage and their R-M systems. The model predicts that sub-populations of specialist phages would evolve.

A second prediction of the model is that the first phage that breaches the restriction barrier of the new host will dominate the population of phages that evolve to specialize on that host, even if the phage has the same affinity for the new host as other phage in the population. This prediction could be tested by beginning the experiment outlined above with a small number of phage with the new methylation pattern that had been marked (e.g. with a small neutral insertion in their genomes) to enable their subsequent identification. The model predicts that many phage in the new specialist population would be descended from the test lineage with the new methylation pattern at the beginning of the experiment.

The third prediction is that the adsorption rate of the first phage to breach the restriction barrier of a new host will only be substantially different from the rest of the population when the mutation rate of the phage is similar to the probability of bypassing its R-M system. The offspring of the first phage to infect a new host can be isolated by plating on that host. Sequencing could reveal if the phages that bypassed the R-M system contained new mutations. The restriction barrier of the new host could be increased or decreased by increasing or decreasing the number of motifs recognized by the R-M system in the genome of the phage (29), or perhaps by increasing or decreasing expression level of the host restriction endonucleases and methyltransferases. Finally, the mutation rate of the phage can be adjusted by growing the phage in the presence of a mutagen. Thus, the prediction could be tested by isolating the first phage to infect a new host for varying rates of mutation and restriction escape.

The model analyzed here did not allow sites recognized by the R-M systems to mutate. However, mutations to remove R-M recognition sites are readily isolated experimentally when phages are not efficiently methylated by host methyltransferases (25). Even when methylation is efficient, as in phage lambda (24), a small cost of plasticity could explain the genomic signature of R-M site avoidance (26). Therefore, an initially plastic host-range phenotype produced by methylation would likely be fixed during evolution (i.e. genetically assimilated (3)). This possibility could be tested during the experiments outlined above by testing for mutations at the R-M recognition sites by sequencing. Experiments to test the role that a cost of plasticity plays on genetic assimilation could also be tested by changing the host’s methylation efficiency. For example, the expression level of the methyltransferase in the host could be increased, which would likely increase the methylation efficiency. Since selection to mutate R-M sites will only operate once a phage infects a new host, I hypothesize that the predictive power of the plastically-produced phenotype will remain high.

## Conclusions

In well-mixed environments, R-M systems provide only temporary protection to bacteria since the first phage to bypass the system produces progeny capable of re-infecting the same host (30). However, the importance of this observation in understanding the role of plasticity in evolution has not been explored. Laboratory evolution experiments cannot determine the events that led to a particular trait in a particular organism in the wild, but they are able to test whether an event *can* cause a particular trait. My results suggest that measuring the predictive power and precedence of plastically-produced phenotypes could elucidate whether they play a causal role on par with mutation during evolution, testing the predictions of plasticity-first evolution.

## Acknowledgements

I thank Daniel R. Matute, Christina L. Burch, Meredith L. Cenzer, Aaron A. Comeault, and Sonia Singhal for helpful comments on the manuscript. I acknowledge the support of the Tri-Institutional Medical Mycology Training Program post-doctoral fellowship (T32-AI052080).

